# Characterization and Organization of Telomeric-Linked Helicase (TLH) Gene Families in *Fusarium oxysporum*

**DOI:** 10.1101/2024.02.27.582403

**Authors:** Sahar Salimi, M. Foad Abdi, Mostafa Rahnama

## Abstract

Telomere-linked RecQ helicase (TLH) genes have been reported in several fungi and a choanoflagellate in the regions adjacent to the terminal telomere repeats. In this study, we explore the Telomere-linked RecQ helicase (TLH) genes in four strains of *Fusarium oxysporum*, offering new insights into their genomic structure, functional motifs, and impact on chromosomal ends. We conducted a comprehensive analysis, comparing the TLH genes of *F. oxysporum* with those previously identified in other organisms and uncovering significant similarities. Through comparative genomics, we identified conserved protein motifs across these genes, including a TLH domain, C_2_H_2_, and RecQ helicase motifs. Our phylogenetic analysis positions the *F. oxysporum* TLH genes in a cluster with other known TLHs, suggesting a shared evolutionary origin. Mutation analysis revealed a relatively low level of deleterious mutations in TLH gene paralogs, with a notable proportion of full-size structures maintained across strains. Analysis of subtelomeric sequences indicates that a region with almost identical sequences flanks the majority of chromosome ends, termed TLH-containing region (TLHcr), across these strains. The presence of TLHcrs at chromosome ends, either as single entities or in arrays, underscores their potential role in telomere function and genome stability. Our findings provide a detailed examination of TLH genes in four strains of *F. oxysporum*, laying the groundwork for future studies on their biological significance and evolutionary history in fungal genomes.

## INTRODUCTION

The *Fusarium oxysporum* species complex is a diverse group of filamentous ascomycete fungi that have an extensive host range from monocotyledonous and dicotyledonous plants (Gordon, 2017; Ma et al., 2013; Michielse, van Wijk, Reijnen, Cornelissen, & Rep, 2009) to immunocompromised humans (O’Donnell et al., 2004) and other mammals (Ortoneda et al., 2004). Although the species as a whole has an extensive host range, individual plant pathogenic isolates of *F. oxysporum* tend to be pathogenic on one or a few species (e.g., only onion, strawberry, or tomato). Isolates with the same host specificity are classified in the same subspecies grouping known as a forma specialis (abbreviated f. sp., plural ff. spp.). For example, pathogenic isolates on strawberries are classified as *F. oxysporum f. sp. fragariae* (Henry et al., 2021; Ma et al., 2013; Michielse, van Wijk, Reijnen, Cornelissen, & Rep, 2009). However, despite the economic significance of these fungi, little is known about their functional diversity that contributes to their pathogenicity and host specificity. One of the significant factors influencing fungal functional diversity is their chromosomal structure, particularly at the end of the chromosomes or telomeres (Langner et al., 2021; Witte et al., 2021). Telomeres are crucial features of linear eukaryotic chromosomes and comprise short, tandem repeats–(TTAGGG)n in most fungi (Blackburn, Greider, & Szostak, 2006; Shay & Wright, 2019). Besides protecting chromosome ends from degradation, telomeres prevent inadvertent end joining by DNA repair mechanisms and extend DNA during replication to overcome sequence loss (Blackburn, Greider, & Szostak, 2006; Vodenicharov & Wellinger, 2006).

Telomeric-linked RecQ-like helicase (TLH) gene families or RecQ-like helicases located at the end of chromosomes or terminal telomeric regions identified in diverse fungi, including *Schizosaccharomyces pombe* (Mandell, Goodrich, Bähler, & Cech, 2005), *Magnaporthe oryzae* (Gao, Khang, Park, Lee, & Kang, 2002; Rehmeyer, Cathryn J., Li, Kusaba, & Farman, 2009) *Ustilago maydis* (Sánchez-Alonso & Guzmán, 1998), *Metarhizium anisopliae* (Inglis, Rigden, Mello, Louis, & Valadares-Inglis, 2005), and the choanoflagellate *Monosiga brevicollis* (Robertson, 2009) underscoring a conserved evolutionary feature across these species. While there is a lack of research explicitly targeting TLH genes many studies concentrate on other RecQ helicases (Paeschke, McDonald, & Zakian, 2010). These helicases belong to superfamily II (SFII) of DNA helicases involved in unwinding DNA duplexes in the 3′ to 5’ direction (Paeschke, McDonald, & Zakian, 2010). This family has been described in various organisms, from prokaryotes to eukaryotes, demonstrating its significant evolutionary conservation (Bachrati & Hickson, 2008; Cobb & Bjergbaek, 2006; Khakhar, Cobb, Bjergbaek, Hickson, & Gasser, 2003; Pérez, Pangilinan, Pisabarro, & Ramírez, 2009; Tuteja & Tuteja, 2004). They play critical roles in

DNA metabolism by preventing illegitimate recombination, repairing stalled replication forks, and initiating homologous recombination, highlighting their essential function across different biological processes (reviewed in (Larsen & Hickson, 2013; Paeschke, McDonald, & Zakian, 2010)). In fission yeast, S. pombe, multiple paralogs of telomere-linked helicase genes (tlh) were found (Mandell, Goodrich, Bähler, & Cech, 2005). Typically silent, these genes become active in the absence of telomerase, particularly during decelerated cell division. The activation of tlh genes significantly enhances the recovery speed from growth arrest in cells lacking telomerase, highlighting the critical role of tlh1 helicase activity in the process of cellular recovery. It is suggested that activating the genes through initiating the Alternative Lengthening of Telomeres (ALT) pathway, a key strategy for maintaining telomeres, contributes to cell recovery (Mandell, Goodrich, Bähler, & Cech, 2005). Similar findings have been reported in the budding yeast *S. cerevisiae*, where Y′ subtelomeric elements encode several helicases called Y′-helicase protein 1 (Y′Help1) expressed under telomerase-deficient cells (Yamada, Hayatsu, Matsuura, & Ishikawa, 1998), underscoring a common evolutionary strategy for telomere maintenance in yeast. In previously described telomere-linked helicase (TLH) proteins, three critical components have been identified: the TLH domain, C_2_H_2_ zinc finger motifs, and RecQ helicase motifs (Rehmeyer, Cathryn J., Li, Kusaba, & Farman, 2009). These components offer significant insights into the operational dynamics of TLH proteins. The C_2_H_2_ zinc finger motifs are essential for the specific DNA-binding capability of these helicases, playing a critical role in the maintenance of telomeres (Brayer & Segal, 2008; Li, Q., Tan, & Wu, 2023). Meanwhile, the RecQ motifs are instrumental in the helicases’ enzymatic activity, enabling them to unwind complex DNA structures found at telomeric ends (Croteau, Popuri, Opresko, & Bohr, 2014; Kaur, Agrawal, & Sengupta, 2021). These functional domains are pivotal for preserving genomic stability, as they help maintain telomere integrity, prevent telomere fusion, and facilitate accurate DNA replication within telomeric regions. This combination of domains within TLH proteins underscores their vital role in safeguarding chromosome ends against degradation and ensuring the cell’s genomic stability.

In this study, we analyzed the subtelomeric regions of four *F. oxysporum* strains, three strains of *F. oxysporum* f. sp. *fragariae* (*Fof*), which causes strawberry Fusarium wilt (Henry et al., 2021), alongside the endophytic strain *Fo*47 (Wang et al., 2020). The investigation led to the discovery of novel telomeric-linked RecQ-like helicase (TLH) genes. Through detailed genomic analysis, we uncovered a specialized region, termed the TLH-containing region (TLHcr), situated near chromosomal termini, enriching our knowledge of telomere architecture in these fungi. In addition, we demonstrated a high level of stability of TLH gene sequences with minimal mutations, alongside mapping the varied presence and configuration of TLHcrs within these strains. These findings underscore the potential role of TLH genes in maintaining genomic integrity and provide insights into the evolutionary dynamics of telomere-associated genes in *F. oxysporum*.

## METODS

### Genome assemblies

Genomic sequences for *F. oxysporum* f. sp. *fragariae* MAFF727510 (*Fof*. MAFF727510), *F. oxysporum* f. sp. *fragariae* BRIP62122 (*Fof*. BRIP62122), *F. oxysporum* f. sp. *fragariae* GL1080 (*Fof*. GL1080), and *F. oxysporum* strain Fo47 were retrieved from the NCBI database using the accession numbers GCA_016164145.2, GCA_016166325.2, GCA_016170085.2, and GCA_013085055.1, respectively.

### Gene prediction and sequence alignments

Genes located adjusted to the chromosome ends were systematically searched for all examined genomes. To identify potential genes within these regions, we employed a variety of gene prediction algorithms, including Augustus (Stanke & Morgenstern, 2005), Maker (Campbell, Holt, Moore, & Yandell, 2014), and ORFfinder (https://www.ncbi.nlm.nih.gov/orffinder/).

Both nucleotide sequence alignments and amino acids alignments were generated with the ClustalOmega algorithm (Sievers & Higgins, 2018) implemented in the Geneious Prime 2022.2.2 package (Biomatters Inc., Boston, MA) using default parameters.

### Analysis of conserved motifs

Conserved motifs and patterns among TLH proteins were identified using the public protein analysis servers Motif Scan (https://myhits.sib.swiss/cgi-bin/motif_scan/) and MOTIF (https://www.genome.jp/tools/motif/), and InterPro (https://www.ebi.ac.uk/interpro/), using default parameters.

### Phylogenetics analysis

RAXMLHPC (Stamatakis, 2014) was used to generate the maximum likelihood tree with the PROTGAMMAAUTO substitution model with 10,000 bootstrapping replications. The best tree was plotted using iTOL v6 (Letunic & Bork, 2021).

### Identification of mutations in TLH gene sequences

A multifasta file was created from a ClustalOmega alignment of TLH gene sequences, and different types of mutations were identified through pairwise comparisons of sequences with the TLH consensus sequence using custom Python script.

## RESULTS

### Identification of telomere-linked helicase (TLH) genes in *F. oxysporum* strains

Analysis of the subtelomeric sequences of *Fof* MAFF727510, *Fof* BRIP62122, *Fof* GL1080, and Fo47 revealed that, within a strain, nearly all chromosomal ends are flanked by regions with almost identical sequences that included a TLH gene. Therefore, we refer to these nearly identical sequences as THL-containing regions (TLHcrs) (see below).

Further analysis of TLHcrs in each *F. oxysporum* strain revealed consensus gene predictions consisting of 4776, 4692, 4889, and 4722 bp ORFs in *Fof*. MAFF727510, *Fof*. BRIP62122, *Fof*. GL1080, and Fo47, respectively (Table 1). In BLAST searches, these ORFs and their derived protein sequences show significant similarities with RecQ helicases, especially those encoded by telomeric genes known as telomere-linked helicases (TLH) in other organisms, including *S. pombe, P. oryzae, U. maydis, M. anisopliae* and *M. brevicollis* (Gao, Khang, Park, Lee, & Kang, 2002; Inglis, Rigden, Mello, Louis, & Valadares-Inglis, 2005; Mandell, Goodrich, Bähler, & Cech, 2005; Rehmeyer, Cathryn J., Li, Kusaba, & Farman, 2009; Robertson, 2009; Sánchez-Alonso & Guzmán, 1998). Combination with further investigation, including comparisons of protein domains and phylogenetic analysis, revealed that these genes in *F. oxysporum* are new members of the TLH genes and RecQ family of DNA helicases (see below).

### *F. oxysporum* TLHs have conserved protein motifs similar with other known TLHs

*F. oxysporum* TLH gene sequences were characterized by generating a TLH consensus sequences for each strain and then comparing the corresponding protein to conceptual TLH proteins reported in other organisms (Figure 1A). The comparison was conducted using four other known fungal TLH proteins and one choanoflagellate TLH protein (Gao, Khang, Park, Lee, & Kang, 2002; Inglis, Rigden, Mello, Louis, & Valadares-Inglis, 2005; Mandell, Goodrich, Bähler, & Cech, 2005; Rehmeyer, Cathryn J., Li, Kusaba, & Farman, 2009; Robertson, 2009; Sánchez-Alonso & Guzmán, 1998). As expected, significant similarity was observed across the RecQ helicase domains, with an average pairwise similarity of 54.6% and an average identity of 27%. High similarity/identity also has been observed in ∼ 250 residues upstream of the helicase domains in the N-terminal region of the TLH proteins, previously referred to as the TLH domain (Rehmeyer, Cathryn J., Li, Kusaba, & Farman, 2009). TLH domains of the ten compared proteins shared 58.5% similarity and 25.7% identity (Figure 1A & B). At the N-terminus ends of all proteins, except for *U. maydis*, two highly conserved C_2_H_2_ Zn^2+^-binding motifs were found. The first C_2_H_2_ motif has the general pattern “Cx_2–3_Cx_8–14_Hx_3–5_H,” except in *M. anisopliae*, in which the second cysteine is replaced with histidine, and the second C_2_H_2_ motif has a Cx_2–5_Cx_11–16_Hx_4–5_H pattern.

**Figure 1.**
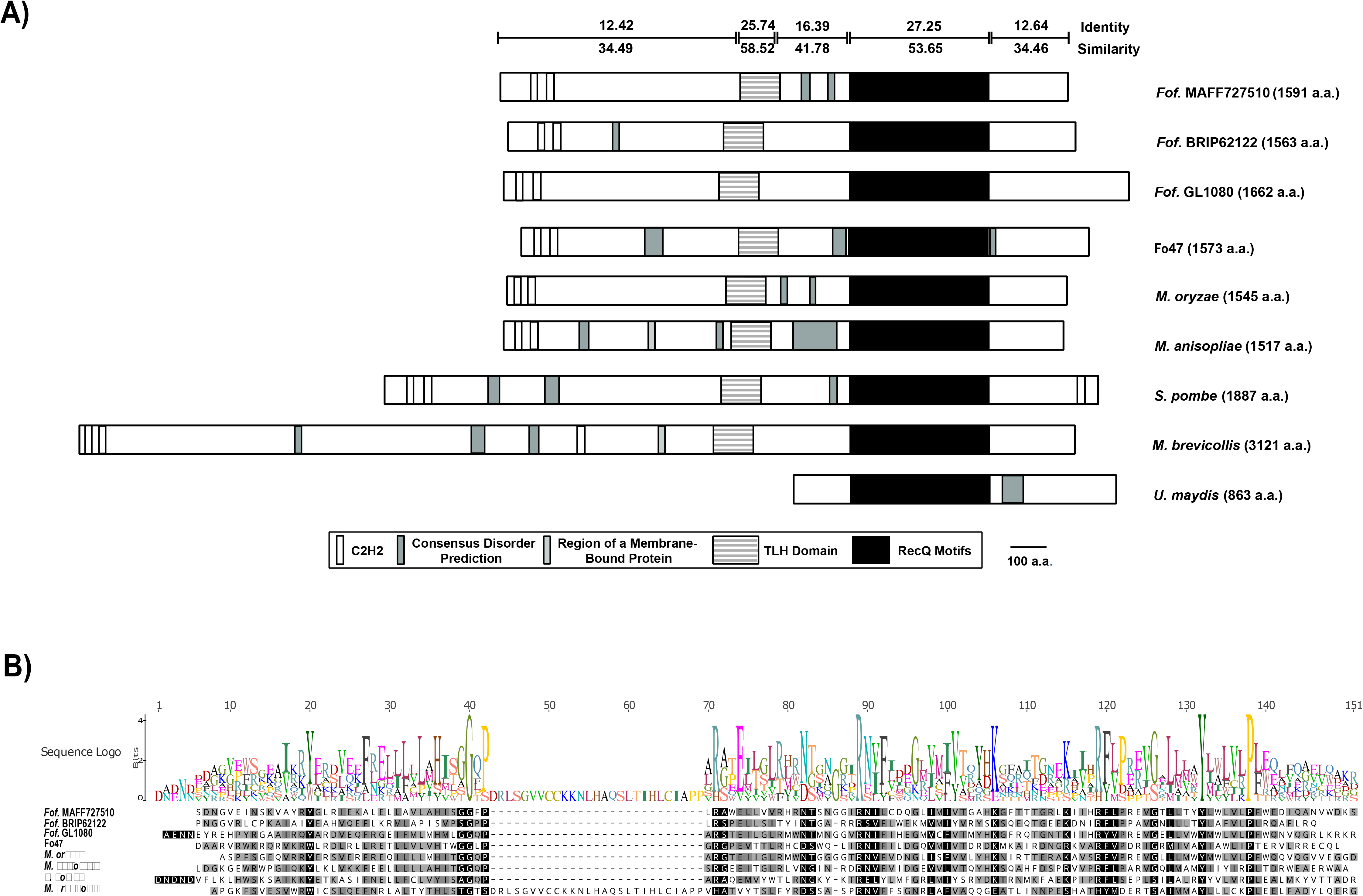
Conserved features in TLH proteins. The TLH protein sequences for *M. oryzae, M. anisopliae, S. pombe, M. brevicollis*, and *U. maydis* were sourced from GenBank (Table S3). **A)** The predicted proteins are aligned based on their RecQ helicase domains. At the top of the figure, the sequence conservation among TLH proteins is represented. Horizontal lines with bars at the end correspond to the regions in the *F. oxysporum* TLH proteins depicted immediately below. Numbers above and below these lines indicate the percentage of identity and similarity, respectively. **B)** Multiple alignment of the TLH domain, with conserved residues highlighted in black and similar residues in gray.

Moreover, three additional C_2_H_2_ motifs were found starting 1758 aa from the N-terminus and 113 aa from the C-terminus of *M. brevicollis* and 74 aa from the C-terminus of *S. pombe* (Figure 1A). We also conducted a phylogenetic analysis of the helicase domain of the four *F. oxysporum* TLHs sequences compared to 18 other selected helicases (Figure 2, Table S3). As shown in the resulting tree topology, the helicases of the four *F. oxysporum* TLHs and the other five known TLHs from *S. pombe, P. oryzae, U. maydis, M. anisopliae*, and *M. brevicollis* formed a diverse cluster from the other helicases examined, supported by 10,000 bootstrap replications (Figure 2).

**Figure 2.**
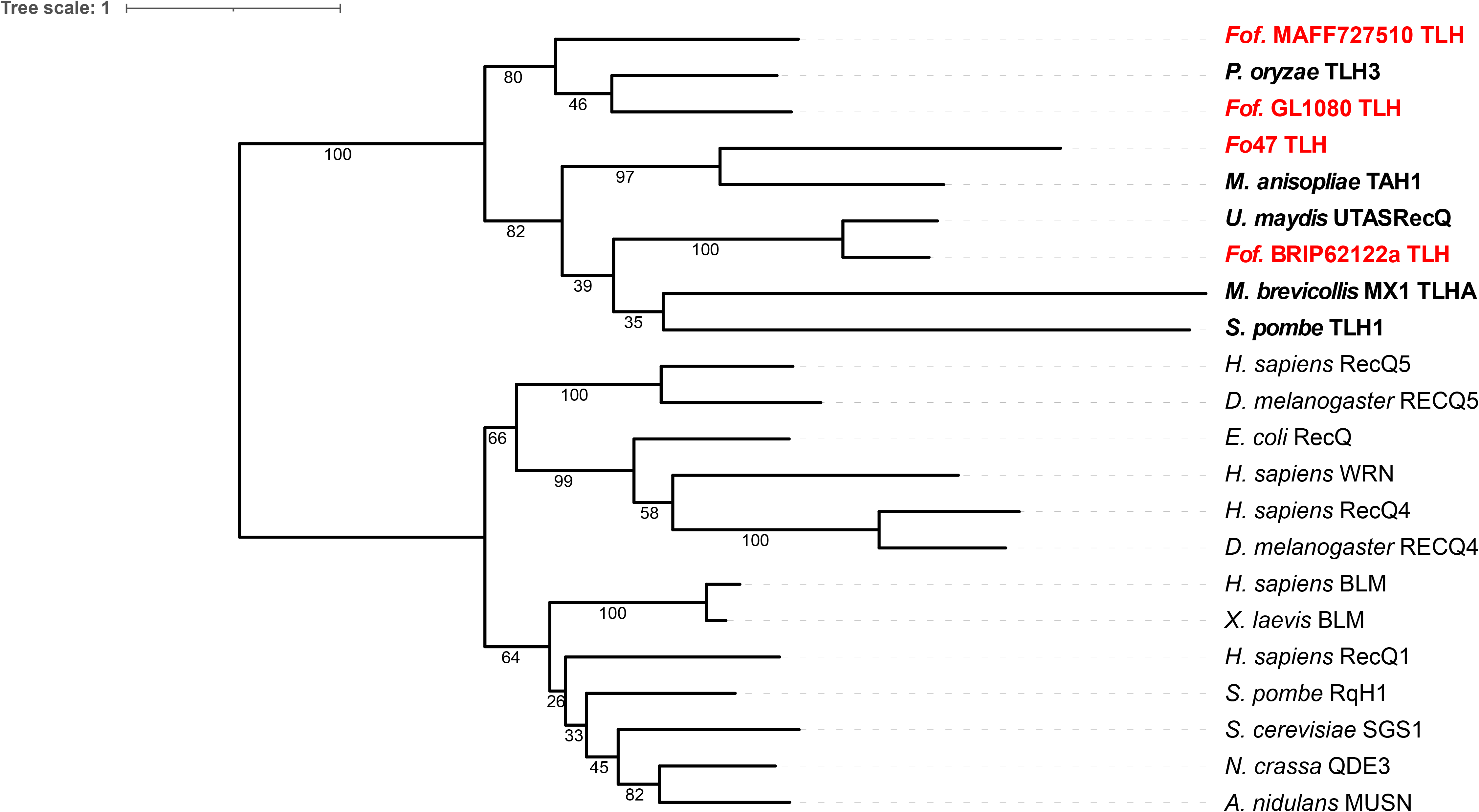
Phylogenetic relationships between *F. oxysporum* TLH genes, other documented TLHs, and various RecQ helicases (Table S3). The tree is based on the central conserved helicase domain of ∽400 amino acids. The *F. oxysporum* proteins are highlighted in red, and other TLH proteins are highlighted in bold. Branch support values represent the percentage of agreement across 10,000 bootstrap replications.

### Mutations in TLH gene paralogs

Examination of TLH ORFs revealed that although most of the paralogs were predicted to encode full-length proteins, some paralogs had nucleotide deletions or insertions that would result in truncated proteins. Each of the four *Fo* strains had at least one mutated TLH paralog: 60% (12 of 20) in *Fof*. MAFF727510, 95% (20 of 21) in *Fof*. BRIP62122, 77% (13 of 17) in *Fof*. GL1080, and 86% (19 of 22) in Fo47, while a small number exhibit mutated (divided) ORFs (Table 1, Figure S2). To identify mutations in the TLH genes in each strain, nucleotide sequences of the TLH genes were aligned using ClustalOmega. This alignment was then compared with its consensus sequence to detect point mutations, nucleotide insertions, and deletions (Table S4).

The results showed an average mutations per gene ranging from 0.29 to 2.5 in *Fof*. BRIP62122 and *Fof*. MAFF727510, respectively (Table 2). Analysis of the alignment in *Fof*. GL1080 revealed a single nucleotide (T) insertion at position 1765 in TLH16. This insertion results in a frameshift, which causes the gene to be divided into two ORFs. In addition, a frameshift was detected in TLH1 and TLH18 due to a deletion in position 4828. Thus, these two paralogs have 158 bp shorter ORFs than the consensus TLH gene in *Fof*. GL1080 (Figure S1A, Table S4). In the *Fof*. BRIP62122, two insertions at position 696 and 1516 divided TLH2 into two ORFs (Figure S1B, Table S4). In the Fo47 strain, three genes are split into two ORFs: TLH5 and TLH14 are divided due to deletions at positions 1704 and 1843, respectively and TLH13-1 because of insertion at position 2483 (Figure S1C, Table S4). Compared to other strains, *Fof*. MAFF727510 TLH genes had a significant higher frequency of mutations (Table 2), resulting in 40% of the ORFs (8 of 20 genes) predicted to yield truncated (Figure S1D, Table S4).

### Most of the chromosome ends are flanked with a TLHcr

Analysis of sequences adjacent to the terminal telomere repeats in the four *F. oxysporum* strains revealed that a majority of them - 83% (20 of 24) in *Fof*. MAFF727510, 91% (20 of 22) in *Fof*. BRIP62122, 79% (22 of 28) in *Fof*. GL1080, and 92% (22 of 24) in Fo47 - are bordered by sequences containing the TLH gene or TLHcr regions (Table S2). TLHcrs exhibit length variability across strains, spanning from 6139bp in *Fof*. GL1080 to 13114bp in *Fof*. MAFF727510 (Figure 3 and Table 1). They are also found in different copy numbers among strains of 43 copies in *Fof*. MAFF727510 to 22 copies in *Fof*. BRIP62122 (Table 1).

**Figure 3.**
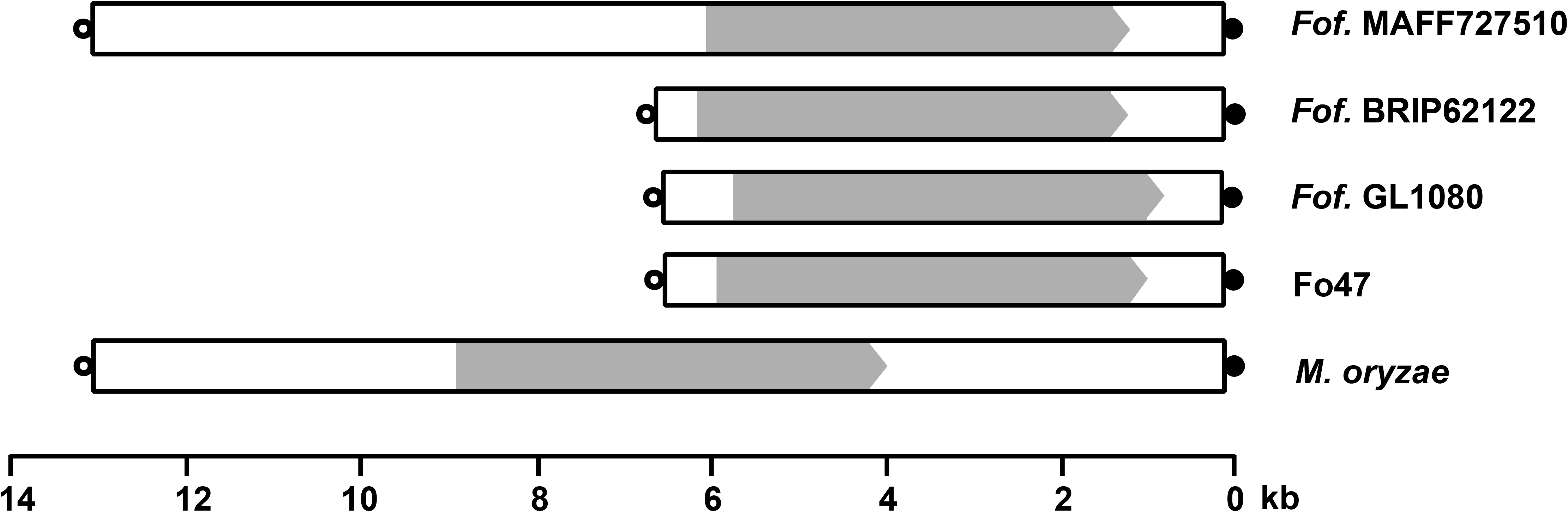
TLHcr (TLH-containing region) structure in *F. oxysporum* strains (top four diagrams) in comparison with *M. oryzae* (bottom diagram). The rectangular boxes depict the TLHcr, and the TLH ORF is indicated with a gray arrow. The filled circles represent terminal telomere repeats, and the empty circles represent interstitial telomere repeat/s.

TLHcrs are positioned at the ends of chromosomes (in full-size or truncated forms) as a single entity or in arrays with a consistent association of telomeric repeats on both sides (Figure 4). The presence of TLHcr arrays is notably higher in the *Fof*. MAFF727510 and *Fof*. GL1080 strains compared to *Fof*. BRIP62122 and Fo47 (Table S2). In the *Fof*. BRIP62122 and Fo47 strains, arrays of two TLHcrs are observed at only two chromosome ends, with the rest featuring a single TLHcr (Table S2). On the other hand, in the *Fof*. MAFF727510, 45% (9 of 20) and in the *Fof*. GL1080 strain, 41% (9 of 22) of chromosome ends associated with TLHcrs are flanked by arrays comprising two or more TLHcrs (Table S2).

**Figure 4.**
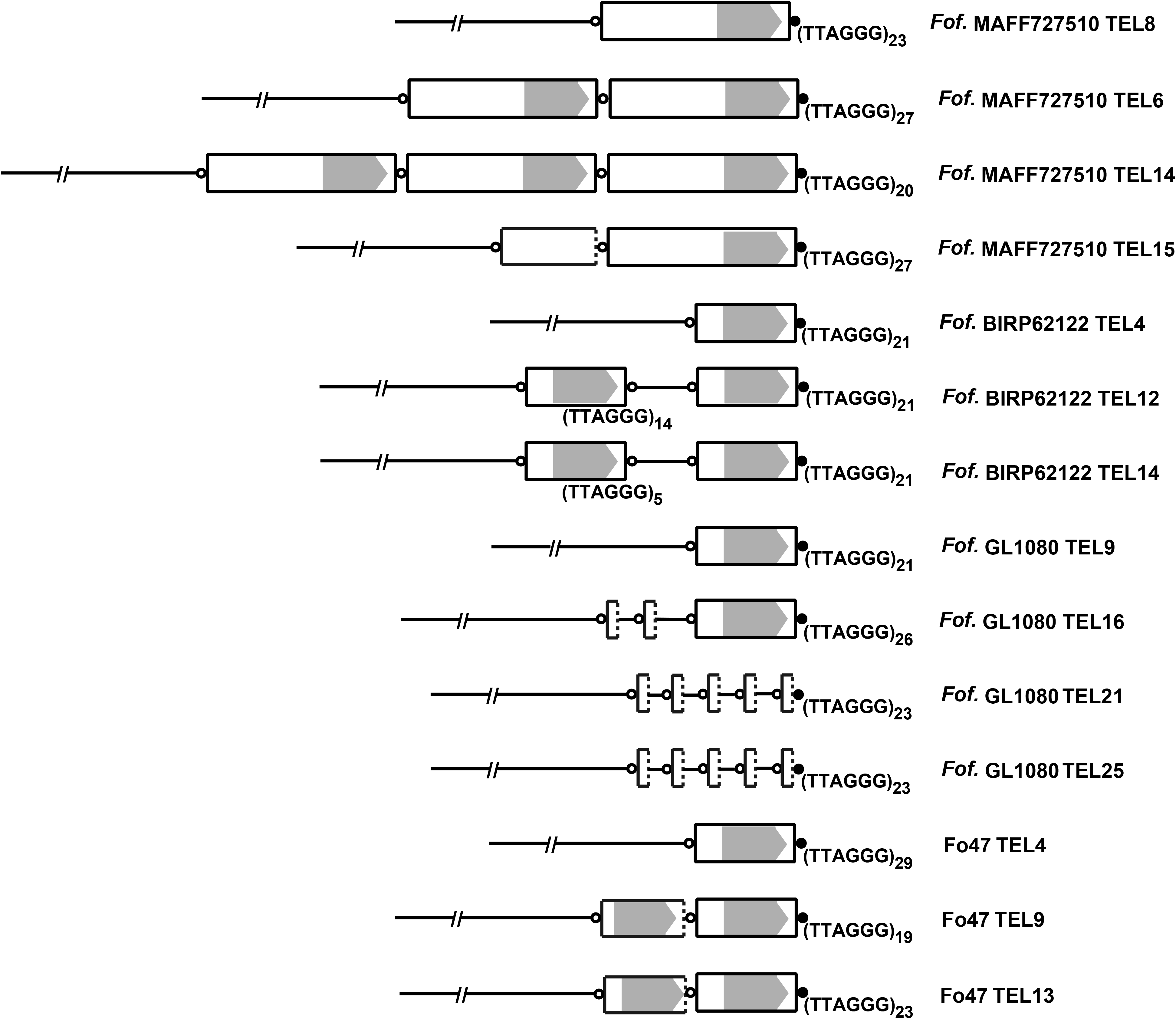
Examples of TLHcrs (TLH-containing regions) organization in the chromosome ends of *F. oxysporum* strains. Rectangular boxes illustrate TLHcrs, with those having dotted angles denoting truncated forms, and TLH ORFs are marked by gray arrows. Solid circles symbolize terminal telomere repeats, whereas open circles indicate interstitial telomere repeats. Each circle denotes a single telomere repeat, except where specific repeat counts are provided beneath them.

## DISCUSSION

Historically, pinpointing telomere-associated genes posed considerable difficulties, mainly because genome assemblies lacked telomeric regions. This challenge was exacerbated in the context of TLH genes, which, as members of multi-gene families situated within expansive, highly duplicated subtelomeric domains, introduced additional layers of complexity (Rehmeyer, Cathryn et al., 2006). However, advancements in sequencing technologies coupled with strides in computational techniques for generating high-quality genome assemblies have markedly improved our understanding of subtelomeric regions, transforming what was once a formidable endeavor into a more manageable task. In this study, by leveraging the existing genome assembly from four *F. oxysporum* strains and refining their subtelomeric regions, we were able to identify TLH genes within these strains. We employed a comparative genomics method to affirm the identification of the TLH ORFs in the *F. oxysporum* strains and compare them with other known TLH genes. The analysis demonstrated notable similarities in the lengths of TLH genes between *F. oxysporum* and those in various other fungi, such as *M. oryzae, M. anisopliae*, and *S. pombe* (Inglis, Rigden, Mello, Louis, & Valadares-Inglis, 2005; Mandell, Goodrich, Bähler, & Cech, 2005; Rehmeyer, Cathryn J., Li, Kusaba, & Farman, 2009; Robertson, 2009). Furthermore, it confirmed the existence of three conserved features within the *F. oxysporum* TLH genes, as previously described in other TLH genes (Rehmeyer, Cathryn J., Li, Kusaba, & Farman, 2009). These include TLH domain, C_2_H_2_, and RecQ motifs, which are positioned with remarkable consistency. In the other previously reported TLH gene in choanoflagellate *M. brevicollis*, despite the TLH ORF being almost twice the length of its fungal counterparts, it exhibits similar motifs and domains with consistent positions (Figure 1A). The observed conserved motifs in *F. oxysporum* and other fungal TLHs, particularly the RecQ helicase and C_2_H_2_, reflect the evolutionary conservation of these proteins. Although the function of the TLH protein is yet to be determined, the broad conservation of its features suggests that they may enhance the functionality or substrate specificity of the TLH protein, thereby augmenting the capabilities of RecQ helicases (Rehmeyer, Cathryn J., Li, Kusaba, & Farman, 2009). The C_2_H_2_ and RecQ domains are vital for helicases’ DNA binding and unwinding functions, respectively (Brayer & Segal, 2008; Paeschke, McDonald, & Zakian, 2010). C_2_H_2_ motifs enable specific interactions with DNA, crucial for telomere maintenance, while RecQ domains are key for enzymatic activity, helping resolve complex DNA structures at telomeres. These domains are central to maintaining genomic stability by preserving telomere integrity, preventing telomere fusion, and ensuring proper DNA replication at telomeric regions (Brayer & Segal, 2008; Croteau, Popuri, Opresko, & Bohr, 2014; Kaur, Agrawal, & Sengupta, 2021; Li, Q., Tan, & Wu, 2023; Paeschke, McDonald, & Zakian, 2010).

The *F. oxysporum* strains vary in number of TLH gene paralogs with the potential capability of encoding full-length proteins — specifically, *Fof*. BRIP62122 and Fo47 have the highest proportion of these paralogs, with 95% and 86% of TLH genes, respectively, while *Fof*. MAFF727510 and *Fof*. GL1080 have the lowest, at 65% and 77%, respectively. In contrast to *F. oxysporum*, in the *M. oryzae* 70-15 strain, a lower number of TLH gene paralogs are predicted to encode full-length proteins (36%, 4 out of 11 genes). When comparing the mutation rates per gene between these species, it becomes evident that *M. oryzae* harbors a substantially higher average number of mutations per gene than *F. oxysporum* — ∼32 in *M. oryzae* to 0.3 to 2.5 *in F. oxysporum* strains. Considering the observed mutation rates in TLH genes, *F. oxysporum* appears to have a higher frequency of TLH genes that yield a functional protein than *M. oryzae*. However, as previously proposed (Rehmeyer, Cathryn J., Li, Kusaba, & Farman, 2009), TLH genes are expected to possess functionality, as non-functional genes would likely be quickly eliminated. Thus, preserving multiple TLH genes across these strains suggests their functional importance and a strong evolutionary selection for their retention.

The observation of TLHcrs at the terminal regions of most chromosomes in *F. oxysporum* strains points to a crucial structural function for these elements. This observation aligns with discoveries in other organisms, including *S. pombe, M. oryzae, M. anisopliae*, and *Pleurotus ostreatus* (Inglis, Rigden, Mello, Louis, & Valadares-Inglis, 2005; Mandell, Goodrich, Bähler, & Cech, 2005; Pérez, Pangilinan, Pisabarro, & Ramírez, 2009; Rehmeyer, Cathryn J., Li, Kusaba, & Farman, 2009), underscoring the broad conservation of TLHcr elements across fungal divisions and classes. Such findings suggest the existence of a potentially universal role for these structures in the stability and maintenance of fungal chromosomes, consistent with the documented conservation of RecQ helicases. Additionally, the variability in TLHcr array configurations observed among *F. oxysporum* strains underscores the genetic intricacies inherent within this species. This spectrum of array variability emphasizes the fluid character of these genomic segments and hints at their significance in ensuring chromosomal stability and managing telomere length. Considering the dynamic nature of subtelomeric regions in other organisms (Brown, Murray, & Verstrepen, 2010; Diotti, Esposito, & Shen, 2021; Jolivet et al., 2019a; Jolivet et al., 2019b; Rahnama et al., 2021), TLHs may be one of the factors contributing to this dynamism. Such insights contribute to our understanding of the molecular mechanisms underpinning genome stability and adaptation in *F. oxysporum*, reflecting broader principles of fungal genetics and chromosomal dynamics. Another notable characteristic of the TLHcr structure is its uniform association with telomeric repeats flanking both sides, irrespective of the elements being full-length or truncated. The interstitial telomere repeats associated with TLHcrs suggest a complex evolutionary history of chromosomal rearrangements, including fusion, duplication, and translocation events (Rahnama et al., 2020; Rahnama et al., 2021). These repeats can serve as sites of chromosomal fragility or recombination hotspots, influencing genome stability (Rahnama et al., 2020)

While the specific functions of TLH genes remain elusive, evidence indicates their crucial role in telomere biology. In both *S. cerevisiae* and *S. pombe*, the typically repressed TLH genes are activated under telomere crisis, suggesting their role in telomere repair (Mandell, Bähler, Volpe, Martienssen, & Cech, 2005; Yamada, Hayatsu, Matsuura, & Ishikawa, 1998). Particularly in *S. pombe*, enhancing the expression of a TLH protein’s C-terminal segment aids in overcoming telomere crisis, indicating TLH proteins’ significant function in telomere maintenance and repair (Mandell, Goodrich, Bähler, & Cech, 2005). The activation of TLH genes, likely modulated by their proximity to telomeres, suggests a model where telomere shortening triggers their expression, possibly through telomere position effect mechanisms, underscoring the adaptive response to maintain telomere integrity (Rehmeyer, Cathryn J., Li, Kusaba, & Farman, 2009).

In conclusion, this study enriches our understanding of the TLH gene’s participation in *F. oxysporum* chromosome structure and draws parallels with other fungal organisms. It highlights the conserved nature of these genes and their potential universal role in chromosomal stability and maintenance across fungi. The findings, in light of the existing literature, open new avenues for exploring TLH’s roles in broader biological contexts and their evolutionary significance. Future research should aim to elucidate the functional implications of these findings, particularly how these genetic elements influence chromosomal stability and adaptation in different *F. oxysporum* groups and other organisms.

## Acknowledgments

The authors thank Li-Jun Ma (University of Massachusetts Amherst) and Robert Proctor (US Department of Agriculture) for the helpful comments, Peter Henry (US Department of Agriculture) for providing PacBio reads, and Michael Renfro (Tennessee Tech University) for technical help. We thank Tennessee Tech University Information Technology Services High-Performance Computing facility for supporting and using their associated resources.

---

**Figure S1.**
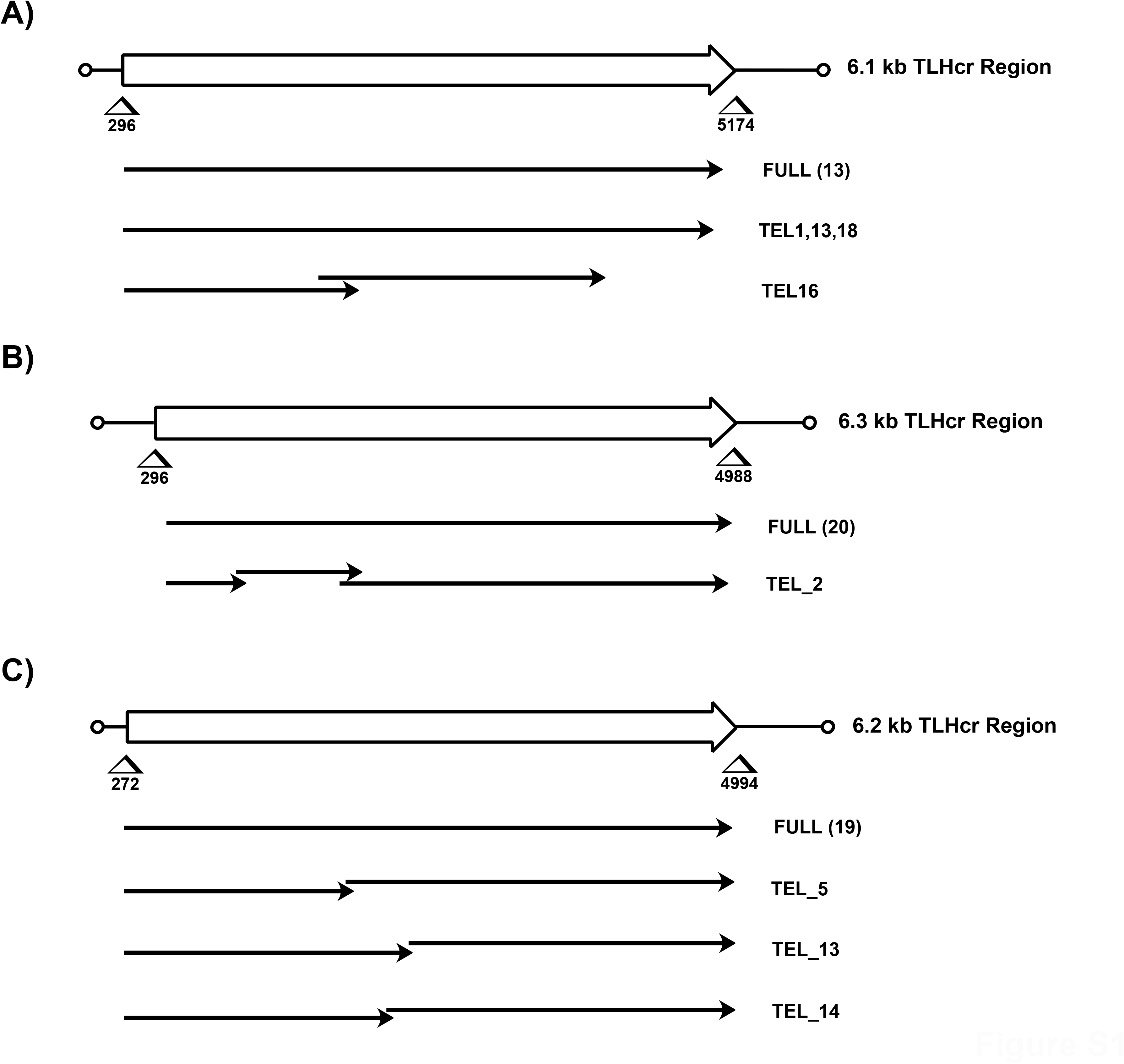
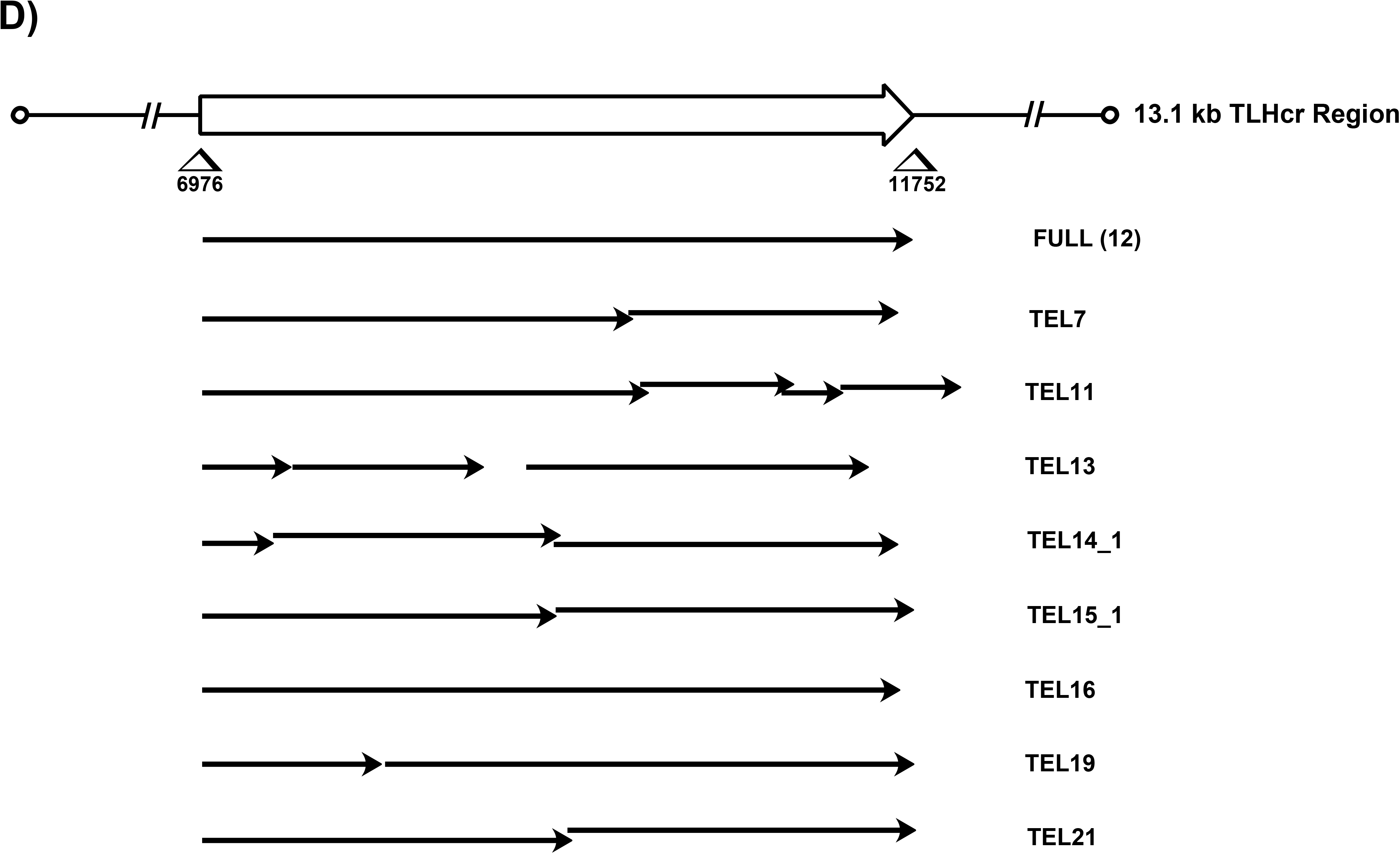
TLH genes organization in *F. oxysporum* strains of **A)** *Fof*. GL1080, **B)** *Fof*. BRIP62122, **C)** Fo47, and **D)** *Fof*. MAFF727510. The boxed arrow at the top of each figure illustrates the predicted TLH ORF overlaid on the TLHcr (TLH-containing regions). The numbers beneath the arrowheads indicate the nucleotide positions within the TLHrc fragment. Solid lines ending in arrowheads depict the predicted genes that coincide with the TLH gene. The initial line signifies a full-size TLH gene, and the number in parentheses shows the count observed. The empty circles represent telomere repeat/s. For additional details see Table S2.

## REFERENCES

Bachrati, C. Z., & Hickson, I. D. (2008). RecQ helicases: Guardian angels of the DNA replication fork. Chromosoma, 117(3), 219–233. doi:10.1007/s00412-007-0142-4

Blackburn, E. H., Greider, C. W., & Szostak, J. W. (2006). Telomeres and telomerase: The path from maize, tetrahymena and yeast to human cancer and aging. Nature Medicine, 12(10), 1133–1138. doi:10.1038/nm1006-1133

Brayer, K. J., & Segal, D. J. (2008). Keep your fingers off my DNA: Protein-protein interactions mediated by C2H2 zinc finger domains. Cell Biochemistry and Biophysics, 50(3), 111–131. doi:10.1007/s12013-008-9008-5

Brown, C. A., Murray, A. W., & Verstrepen, K. J. (2010). Rapid expansion and functional divergence of subtelomeric gene families in yeasts. Current Biology : CB, 20(10), 895–903. doi:10.1016/j.cub.2010.04.027

Campbell, M. S., Holt, C., Moore, B., & Yandell, M. (2014). Genome annotation and curation using MAKER and MAKER-P. Current Protocols in Bioinformatics, 48, 4.11.1–4.11.39. doi:10.1002/0471250953.bi0411s48

Cobb, J. A., & Bjergbaek, L. (2006). RecQ helicases: Lessons from model organisms. Nucleic Acids Research, 34(15), 4106–4114. doi:10.1093/nar/gkl557

Croteau, D. L., Popuri, V., Opresko, P. L., & Bohr, V. A. (2014). Human RecQ helicases in DNA repair, recombination, and replication. Annual Review of Biochemistry, 83, 519–552. doi:10.1146/annurev-biochem-060713-035428

Diotti, R., Esposito, M., & Shen, C. H. (2021). Telomeric and sub-telomeric structure and implications in fungal opportunistic pathogens. Microorganisms, 9(7), 1405. doi:10.3390/microorganisms9071405

Gao, W., Khang, C. H., Park, S., Lee, Y., & Kang, S. (2002). Evolution and organization of a highly dynamic, subtelomeric helicase gene family in the rice blast fungus magnaporthe grisea. Genetics, 162(1), 103–112. Retrieved from https://www.ncbi.nlm.nih.gov/pmc/articles/PMC1462230/

Gordon, T. R. (2017). Fusarium oxysporum and the fusarium wilt syndrome. Annual Review of Phytopathology, 55, 23–39. doi:10.1146/annurev-phyto-080615-095919

Henry, P. M., Pincot, D. D. A., Jenner, B. N., Borrero, C., Aviles, M., Nam, M., … Gordon, T. R. (2021). Horizontal chromosome transfer and independent evolution drive diversification in fusarium oxysporum f. sp. fragariae. The New Phytologist, 230(1), 327–340. doi:10.1111/nph.17141

Inglis, P. W., Rigden, D. J., Mello, L. V., Louis, E. J., & Valadares-Inglis, M. C. (2005). Monomorphic subtelomeric DNA in the filamentous fungus, metarhizium anisopliae,contains a RecQ helicase-like gene. Molecular Genetics and Genomics: MGG, 274(1), 79–90. doi:10.1007/s00438-005-1154-5

Jolivet, P., Serhal, K., Graf, M., Eberhard, S., Xu, Z., Luke, B., & Teixeira, M. T. (2019a). A subtelomeric region affects telomerase-negative replicative senescence in saccharomyces cerevisiae. Scientific Reports, 9(1), 1845. doi:10.1038/s41598-018-38000-9

Jolivet, P., Serhal, K., Graf, M., Eberhard, S., Xu, Z., Luke, B., & Teixeira, M. T. (2019b). A subtelomeric region affects telomerase-negative replicative senescence in saccharomyces cerevisiae. Scientific Reports, 9(1), 1845. doi:10.1038/s41598-018-38000-9

Kaur, E., Agrawal, R., & Sengupta, S. (2021). Functions of BLM helicase in cells: Is it acting like a double-edged sword? Frontiers in Genetics, 12, 634789. doi:10.3389/fgene.2021.634789

Khakhar, R. R., Cobb, J. A., Bjergbaek, L., Hickson, I. D., & Gasser, S. M. (2003). RecQ helicases: Multiple roles in genome maintenance. Trends in Cell Biology, 13(9), 493–501. doi:10.1016/s0962-8924(03)00171-5

Langner, T., Harant, A., Gomez-Luciano, L. B., Shrestha, R. K., Malmgren, A., Latorre, S. M., … Kamoun, S. (2021). Genomic rearrangements generate hypervariable mini-chromosomes in host-specific isolates of the blast fungus. PLOS Genetics, 17(2), e1009386. doi:10.1371/journal.pgen.1009386

Larsen, N. B., & Hickson, I. D. (2013). RecQ helicases: Conserved guardians of genomic integrity. Advances in Experimental Medicine and Biology, 767, 161–184. doi:10.1007/978-1-4614-5037-5_8

Letunic, I., & Bork, P. (2021). Interactive tree of life (iTOL) v5: An online tool for phylogenetic tree display and annotation. Nucleic Acids Research, 49(W1), W293–W296. doi:10.1093/nar/gkab301

Li, H. (2018). Minimap2: Pairwise alignment for nucleotide sequences. Bioinformatics (Oxford, England), 34(18), 3094–3100. doi:10.1093/bioinformatics/bty191

Li, Q., Tan, G., & Wu, F. (2023). The functions and roles of C2H2 zinc finger proteins in hepatocellular carcinoma. Frontiers in Physiology, 14, 1129889. doi:10.3389/fphys.2023.1129889

Ma, L., Geiser, D. M., Proctor, R. H., Rooney, A. P., O’Donnell, K., Trail, F., … Kazan, K. (2013). Fusarium pathogenomics. Annual Review of Microbiology, 67, 399–416. doi:10.1146/annurev-micro-092412-155650

Mandell, J. G., Goodrich, K. J., Bähler, J., & Cech, T. R. (2005). Expression of a RecQ helicase homolog affects progression through crisis in fission yeast lacking telomerase. The Journal of Biological Chemistry, 280(7), 5249–5257. doi:10.1074/jbc.M412756200

Mandell, J. G., Bähler, J., Volpe, T. A., Martienssen, R. A., & Cech, T. R. (2005). Global expression changes resulting from loss of telomeric DNA in fission yeast. Genome Biology, 6(1), R1. doi:10.1186/gb-2004-6-1-r1

Michielse, C. B., van Wijk, R., Reijnen, L., Cornelissen, B. J. C., & Rep, M. (2009). Insight into the molecular requirements for pathogenicity of fusarium oxysporum f. sp. lycopersici through large-scale insertional mutagenesis. Genome Biology, 10(1), R4. doi:10.1186/gb-2009-10-1-r4

O’Donnell, K., Sutton, D. A., Rinaldi, M. G., Magnon, K. C., Cox, P. A., Revankar, S. G., … Robinson, J. S. (2004). Genetic diversity of human pathogenic members of the fusarium oxysporum complex inferred from multilocus DNA sequence data and amplified fragment length polymorphism analyses: Evidence for the recent dispersion of a geographically widespread clonal lineage and nosocomial origin. Journal of Clinical Microbiology, 42(11), 5109–5120. doi:10.1128/JCM.42.11.5109-5120.2004

Ortoneda, M., Guarro, J., Madrid, M. P., Caracuel, Z., Roncero, M. I. G., Mayayo, E., & Di Pietro, A. (2004). Fusarium oxysporum as a multihost model for the genetic dissection of fungal virulence in plants and mammals. Infection and Immunity, 72(3), 1760–1766. doi:10.1128/IAI.72.3.1760-1766.2004

Paeschke, K., McDonald, K. R., & Zakian, V. A. (2010). Telomeres: Structures in need of unwinding. FEBS Letters, 584(17), 3760–3772. doi:10.1016/j.febslet.2010.07.007

Pérez, G., Pangilinan, J., Pisabarro, A. G., & Ramírez, L. (2009). Telomere organization in the ligninolytic basidiomycete pleurotus ostreatus. Applied and Environmental Microbiology, 75(5), 1427–1436. doi:10.1128/AEM.01889-08

Rahnama, M., Novikova, O., Starnes, J. H., Zhang, S., Chen, L., & Farman, M. L. (2020). Transposon-mediated telomere destabilization: A driver of genome evolution in the blast fungus. Nucleic Acids Research, 48(13), 7197–7217. doi:10.1093/nar/gkaa287

Rahnama, M., Wang, B., Dostart, J., Novikova, O., Yackzan, D., Yackzan, A., … Farman, M. L. (2021). Telomere roles in fungal genome evolution and adaptation. Frontiers in Genetics, 12, 676751. doi:10.3389/fgene.2021.676751

Rehmeyer, C. J., Li, W., Kusaba, M., & Farman, M. L. (2009). The telomere-linked helicase (TLH) gene family in magnaporthe oryzae: Revised gene structure reveals a novel TLH-specific protein motif. Current Genetics, 55(3), 253–262. doi:10.1007/s00294-009-0240-3

Rehmeyer, C., Li, W., Kusaba, M., Kim, Y., Brown, D., Staben, C., … Farman, M. (2006). Organization of chromosome ends in the rice blast fungus, magnaporthe oryzae. Nucleic Acids Research, 34(17), 4685–4701. doi:10.1093/nar/gkl588

Robertson, H. M. (2009). The choanoflagellate monosiga brevicollis karyotype revealed by the genome sequence: Telomere-linked helicase genes resemble those of some fungi. Chromosome Research: An International Journal on the Molecular, Supramolecular and Evolutionary Aspects of Chromosome Biology, 17(7), 873–882. doi:10.1007/s10577-009-9078-2

Sánchez-Alonso, P., & Guzmán, P. (1998). Organization of chromosome ends in ustilago maydis. RecQ-like helicase motifs at telomeric regions. Genetics, 148(3), 1043–1054. doi:10.1093/genetics/148.3.1043

Shay, J. W., & Wright, W. E. (2019). Telomeres and telomerase: Three decades of progress. Nature Reviews. Genetics, 20(5), 299–309. doi:10.1038/s41576-019-0099-1

Sievers, F., & Higgins, D. G. (2018). Clustal omega for making accurate alignments of many protein sequences. Protein Science : A Publication of the Protein Society, 27(1), 135–145. doi:10.1002/pro.3290

Stamatakis, A. (2014). RAxML version 8: A tool for phylogenetic analysis and post-analysis of large phylogenies. Bioinformatics (Oxford, England), 30(9), 1312–1313. doi:10.1093/bioinformatics/btu033

Stanke, M., & Morgenstern, B. (2005). AUGUSTUS: A web server for gene prediction in eukaryotes that allows user-defined constraints. Nucleic Acids Research, 33(Web Server issue), W465–W467. doi:10.1093/nar/gki458

Thorvaldsdóttir, H., Robinson, J. T., & Mesirov, J. P. (2013). Integrative genomics viewer (IGV): High-performance genomics data visualization and exploration. Briefings in Bioinformatics, 14(2), 178–192. doi:10.1093/bib/bbs017

Tuteja, N., & Tuteja, R. (2004). Prokaryotic and eukaryotic DNA helicases. essential molecular motor proteins for cellular machinery. European Journal of Biochemistry, 271(10), 1835–1848. doi:10.1111/j.1432-1033.2004.04093.x

Vodenicharov, M. D., & Wellinger, R. J. (2006). DNA degradation at unprotected telomeres in yeast is regulated by the CDK1 (Cdc28/clb) cell-cycle kinase. Molecular Cell, 24(1), 127–137. doi:10.1016/j.molcel.2006.07.035

Wang, B., Yu, H., Jia, Y., Dong, Q., Steinberg, C., Alabouvette, C., … Guo, L. (2020). Chromosome-scale genome assembly of fusarium oxysporum strain Fo47, a fungal endophyte and biocontrol agent. Molecular Plant-Microbe Interactions: MPMI, 33(9), 1108–1111. doi:10.1094/MPMI-05-20-0116-A

Witte, T. E., Harris, L. J., Nguyen, H. D. T., Hermans, A., Johnston, A., Sproule, A., … Overy, D. P. (2021). Apicidin biosynthesis is linked to accessory chromosomes in fusarium poae isolates. BMC Genomics, 22(1), 591. doi:10.1186/s12864-021-07617-y

Yamada, M., Hayatsu, N., Matsuura, A., & Ishikawa, F. (1998). Y’-Help1, a DNA helicase encoded by the yeast subtelomeric Y’ element, is induced in survivors defective for telomerase. The Journal of Biological Chemistry, 273(50), 33360–33366. doi:10.1074/jbc.273.50.33360

